# Mapping the daily rhythmic transcriptome in the diabetic retina

**DOI:** 10.1101/2023.05.27.542572

**Authors:** Ryan P. Silk, Hanagh R. Winter, Ouria Dkhissi -Benyahya, Carmella Evans-Molina, Alan W. Stitt, Vijay K. Tiwari, David A. Simpson, Eleni Beli

## Abstract

Retinal function shows marked changes from day to night. Yet, clinical diagnosis, treatments, and experimental sampling occur during the day, leaving a significant gap in our understanding of the pathobiology occurring at night. While there is evidence that diabetes disrupts the circadian system that optimizes our physiology to the environmental light/dark cycle, the impact of such disruption is not well understood. This study investigates whether diabetes affects the retina’s daily rhythm of gene expression to understand the pathobiology of diabetic retinopathy. Ins2^Akita/J^ mice, a model of type 1 diabetes, were kept under a standard 12h:12h light/dark cycle until four months of age. Non-diabetic littermates were used as controls. Bulk mRNA sequencing was conducted in retinas collected every 4 hours throughout the 24 hr light/dark cycle. Computational approaches were used to detect rhythmicity, predict acrophase, identify differential rhythmic patterns, analyze phase set enrichment, and predict upstream regulators. The retinal transcriptome exhibited a tightly regulated rhythmic expression with a clear 12-hr axis of transcriptional rush, peaking at midday and midnight. The functions of day-peaking genes were enriched for DNA repair, RNA splicing, and ribosomal protein synthesis, whereas night-peaking genes were enriched for metabolic processes and growth factor signaling. Although the 12-hr transcriptional axis is retained in the diabetic retina, it was phase advanced by approximately 1-3 hours with a wider distribution. Upstream regulator analysis for the genes that showed phase shifts identified oxygen sensing mechanisms and HIF1alpha as regulators, but not the circadian clock, which remained in phase to the light/dark cycle. We propose a model in which early in diabetes, the retina experiences a jet lag caused by the entrained circadian clock and its output being in one phase and metabolic pathways related to neuronal dysfunction and hypoxia driving advancement of gene expression to a different phase. Further studies are now required to evaluate the chronic implications of such internal jet lag for development of diabetic retinopathy.

## Introduction

With the inexorable increase in the incidence of diabetes, it is estimated that by 2040 around 224 million people will have some form of diabetic retinopathy (DR), with approximately 70 million suffering severe vision loss (1). Current treatments are administered only at the late stages of the disease and do not always prevent vision loss, carry side effects, and place a substantial economic burden on healthcare systems worldwide. Early diagnosis, prevention strategies, and new therapies are urgently needed. In this respect, understanding early pathobiology and how environmental inputs impact disease progression can provide the basis for designing prevention strategies by modifying these inputs.

DR pathogenesis is complex and incompletely elucidated. Hyperglycemia drives early-stage retinal gliosis, neuroinflammation, and vascular pathology, which progresses to ischemia and the sight-threatening stages typified by neuronal depletion and loss of retinal integrity due to edema and neovascularization. At the early stages, most clinical efforts focus on managing glucose levels and blood pressure with proportionately less provision for lifestyle or environmental changes that can benefit patients.

While it is well established that circadian disruption predisposes to diabetes (2-5), disease-induced disruption of the circadian system emerges as an important factor for prognosis of disease management and outcomes. Specifically, circadian disruption in diabetes seems to impact peripheral clocks more than the master clock located in the suprachiasmatic nucleus (4, 6) leading to a misalignment between the circadian clocks throughout the body. Additionally, a metabolic jet-lag which refers to a state of phase shift in circadian profiles of metabolic or endocrine pathways was described for people with aberrant eating behaviors (7) but it can also occur in diabetes.

The circadian clock, a molecular mechanism, adapts multiple aspects of our physiology to the daily light cycle. It comprises a group of proteins operating in a translational transcriptional feedback loop (TTFL) that takes approximately 24 hours to complete. By regulating the rhythmic transcription of genes at the cellular level, this system coordinates multiple aspects of our physiology to adapt to the daily light/dark cycles. As in all our tissues, circadian clocks operate in the retina (8-10) and prepare it for the metabolic and functional changes of the day-night cycle. There is clear evidence that diabetes affects retinal circadian clock rhythms and alters circadian gene expression in the retina in both type 1 and type 2 diabetic models (6, 11-14). Importantly, this disruption occurs before the appearance of morphological changes associated with a diagnosis of DR (6).

While these studies indicate clearly that diabetes disrupts how this molecular mechanism works, light can entrain the clock daily. At least one study, indicates that light-induced circadian gene expression in the retina is also affected in diabetes (15) due to reduced c-fos and dopamine release upon light stimulation. Additional studies indicate that under physiological light/dark conditions the daily rhythms of activity, blood pressure, and temperature (16, 17), the rhythms of glucose and insulin sensitivity (18, 19) circulating immune cells (20), metabolites (21, 22) and the microbiota (21, 23) are impaired by diabetes.

This study was designed to map the gene expression changes in control and diabetic retinas under physiological light/dark conditions that correspond to what diabetic patients experience in their daily life. We hypothesized that diabetes would influence the global daily rhythmic transcriptome in the retina. We collected retinas from four-month-old Ins2Akita mice at different time points in the day/light cycle and performed global mRNA sequencing. We show that very early in diabetes and before the appearance of DR pathology, a significant misalignment of the retinal rhythmic outputs occurs, possibly leading to the manifestation of DR. We confirm that functional compartmentalization of transcriptional rush-times takes place, with a 12hr phase difference in day and night are present in the retina. We finally show that diabetes results in phase shifts and amplitude changes, and we determined the most significant pathways impacted and their possible regulators. Finally, we propose that diabetes leads to an internal desynchrony between metabolic and circadian-led rhythms that could contribute to the pathogenesis of DR.

## Methods and Materials

### Ethics Statement

Animal studies were carried out at the institutional animal care facilities at the Indiana University School of Medicine per institutional and national guidelines for the care and use of laboratory animals (IACUC #10604 and #11167)

### Mouse Model

The C57BL/6-Ins2^Akita^/J (Ins2^Akita^) mouse model of T1D was selected in this study. Heterozygous male Ins2^Akita^ mice develop severe hyperglycemia by 4.5 weeks resulting from insulin misfolding and beta cell destruction. The retinas of these mice exhibit vascular, neural, and glial abnormalities consistent with clinical observations of early diabetic retinopathy (24). Mice were bred at Indiana University according to approved IACUC. Male Ins2^Akita^ heterozygous mice were bred with female C57Bl6 mice, and heterozygous male Ins2^Akita^ exhibited consistent hyperglycemia (>250mg/dL), while littermate males without phenotype considered controls. Mice were kept at constant temperature and humidity under a 12h:12h Light/Dark (12L/12D) schedule from birth, with *ad libitum* access to food and water, and when they reached four months of age were euthanized for sample collection. This age was chosen as a narrow window where mice have established diabetes but still do not develop overt DR, which occurs after six months of age (24). Sampling was performed at different times of the day/night cycle.

### Sample Collection

Mice were selected randomly at four months of age and euthanized at one of six time-points over a 24hour light/dark cycle (ZT: 1, 5, 9, 13, 17, 21 where ZT0 is indicative of lights on and ZT12 lights off), with 4-5 biological replicates for each time-point **(S.Fig. 1)**. Terminal perfusion was conducted using cold phosphate-buffered saline (PBS) solution (without Ca^2+^ and Mg^2+^), eyes enucleated, and retinas were dissected under a microscope. For the night points (ZT12-ZT24), all processes were performed under dim red light. Following isolation, retinas were immediately frozen in liquid nitrogen and stored at -80°C. Processing of samples took ∼10 minutes for each sample and ∼1 hour for each ZT group, starting 30 minutes before the time-point and finishing 30 min after. Frozen samples were shipped to Queen’s University Belfast for preparation and sequencing.

### Sample Preparation and Sequencing

For individual retinal samples, total RNA isolation was performed using the Qiagen RNeasy Mini Kit. Complementary DNA (cDNA) libraries were constructed using the KAPA RNA HyperPrep Kit with RiboErase (HMR) amplification and quality control procedures conducted for individual cDNA libraries. Generated Libraries were quantified using Kapa Quantification, normalized, and pooled in equimolar amounts, and quality control was performed. RNA seq was conducted in the Queens’ University Belfast Genomics Core Technology Unit (https://www.qub.ac.uk/sites/core-technology-units/Genomics/) using 100-cycle Illumina NovaSeq 6000 S1 paired-end (2 x 50bp) sequencing, which produced an average read depth of approximately 27.5 million reads per sample.

### Read Processing, Mapping, and Quantification

Raw sequence reads for diabetic and healthy control retinas were initially inspected for quality using FastQC (25). Raw sequence reads were mapped to the *Mus musculus* reference genome (GRCm38.p5) and annotated using Gencode (M15) with the STAR aligner program (v2.7.4a)(26). HTSeq-count (v0.11.1)(27) was used to generate the raw counts for each sample, and the resultant read count matrix was normalized together using DESeq2 (v1.26.0)(28). To increase statistical power for identifying truly cycling gene features with extremely low read counts (29), following normalization and visual inspection of the data, genes with a normalized median read count >4 across all samples were considered expressed and included for further analysis. The average read count was 31,758,612 reads and annotated genes expressed in control and diabetic retinas can be available upon request.

### Identification of Cycling Genes

Genes displaying an approximately 24-hour rhythm were identified using the nonparametric rhythmicity detection algorithm empirical JTK_CYCLE with asymmetry search (empJTK) (30). Briefly, empJTK is an advancement on the well-accepted JTK_CYCLE rhythmicity detection method, which allows the identification of transcripts displaying asymmetric waveforms and is considered one of the most robust methods for identifying periodic patterns in gene expression data (31). Normalized diabetic and control read count matrices were formatted as a single cycle with 4-5 biological replicates for each timepoint, and both phase and asymmetry searches were set to 4-hour intervals ranging from ZT0 to ZT20. Significantly cycling genes were defined as those with an empJTK empirically calculated p-value (empP) <0.05 and an empJTK predicted min/max fold-change of >1.25. empJTK outputs for all datasets analyzed are available upon request. The Metascape online tool (32) was additionally used to compare multiple gene lists.

### Acrophase Prediction

On account of empJTK is only effective at predicting acrophases no more densely than the resolution of the data, harmonic regression was subsequently used to accurately predict the acrophases of significantly cycling genes using the Harmonic Regression R package (v.1.0) with a fixed period of 24 hrs and normalization set to false. Radian acrophases (*φ*) were then converted to hours (*φ′*) using the equation *φ′ =(− P/2π)φ*, where *P* is the pre-set period of rhythmicity (24 hrs) (33).

### Functional Annotation and Phase Set Enrichment Analysis

To analyze the functions of cycling genes while considering the acrophase of the transcript, a DO and KEGG phase set enrichment analysis (PSEA) (34) was carried out using gene sets from the Molecular Signatures Database (MSigDB)(35). Any GO or KEGG term with a BH-corrected p-value < 0.01 was significantly enriched. Harmonic regression predicted acrophases were rounded to the nearest hour. Testing was conducted on a uniform background using the Kuiper test. If sets contained less than ten transcripts, they were excluded from the analysis.

### Differential Rhythmicity Detection

Differential gene expression at each time points and over all time points was performed using DESeq2 (28). Differentially expressed transcripts were defined as those with a Benjamini-Hochberg adjusted p-value < 0.05. Data are shown in **S. Table 5**. To test for rhythmicity changes in amplitude and phase of genes that had been identified as significantly cycling in either diabetic or control retinas, the DODR R package (v0.99.2) was used, (36). Normalized read counts were used with the robust DODR method with a fixed period of 24 hrs. The resultant DODR p-values were Benjamini-Hochberg (BH) adjusted for multiple testing, and an adjusted p-value <0.05 was used to define differential rhythmicity. Differentially rhythmic transcripts were then further subdivided into those displaying a phase shift when the difference in acrophase between diabetic and control > 1 hour and those displaying a substantially altered amplitude with absolute log_2_ amplitude fold-change >0.32.

### Qiagen Ingenuity Pathway Analysis (IPA)

The list of differential rhythmic genes was used for IPA Core Analysis and canonical pathway analysis. The graphical summary of the results was used to understand how the pathways are inter-related to one another. Lists of genes that showed significant phase shifts or amplitude changes were used for upstream regulator analysis tin order to understand the possible upstream factors that drive either of those changes.

### Data Availability

Data for RNA-seq are deposited on the GEO repository (GSE233440). All other data supporting the findings of this study are included in the manuscript and its supplementary files or available from the corresponding authors on request.

## Results

### Daily expression of clock genes in the diabetic retina

A list of 15 known clock genes from the primary (*Arntl, Clock, Npas2, Per1, Per2, Per3, Cry1, Cry2*), secondary (*Nrd1d1, Nr1d2, Rora, Rorb, Rorc*), and accessory TTFL loops (*Nfil3, Dbp*) **(Fig. 1A)** were selected to investigate the effect of diabetes on their rhythmic expression. Seven clock genes were identified as rhythmic in the retinas in the control group (*Arntl, Npas2, Per2, Rorb, Rorc, Dbp, Nfil3*) and eight in the retinas in the diabetic group (*Npas2, Per1, Per2, Rora, Rorb, Rorc, Dbp, Nfil3*) **(Fig. 1B)**. Overall, these data showed limited effects of diabetes on clock gene expression in the retina. Yet, *Arntl1* was rhythmic only in controls, while *Per1*, and *Rora* only in diabetic retinas, while *Per2* and *Rorb* had slightly increased amplitude, notably, the phase of most clock genes in the retina did not change, expect for *Rorb* and *Npas*. We must note that the retinal clock is entrained by the light/dark cycle so differences in the phase under these conditions were not expected. To confirm this, we then analyzed several clock-controlled genes (*Adcy1, Drd4, Nr2e3, Cys1, Plekbh1, Usp2*) (**S.Fig. 2**) that also showed amplitude changes but no phase changes, indicating that the clock is entrained and with the 12L:12D cycle at this stage of diabetes.

**Figure 1.**
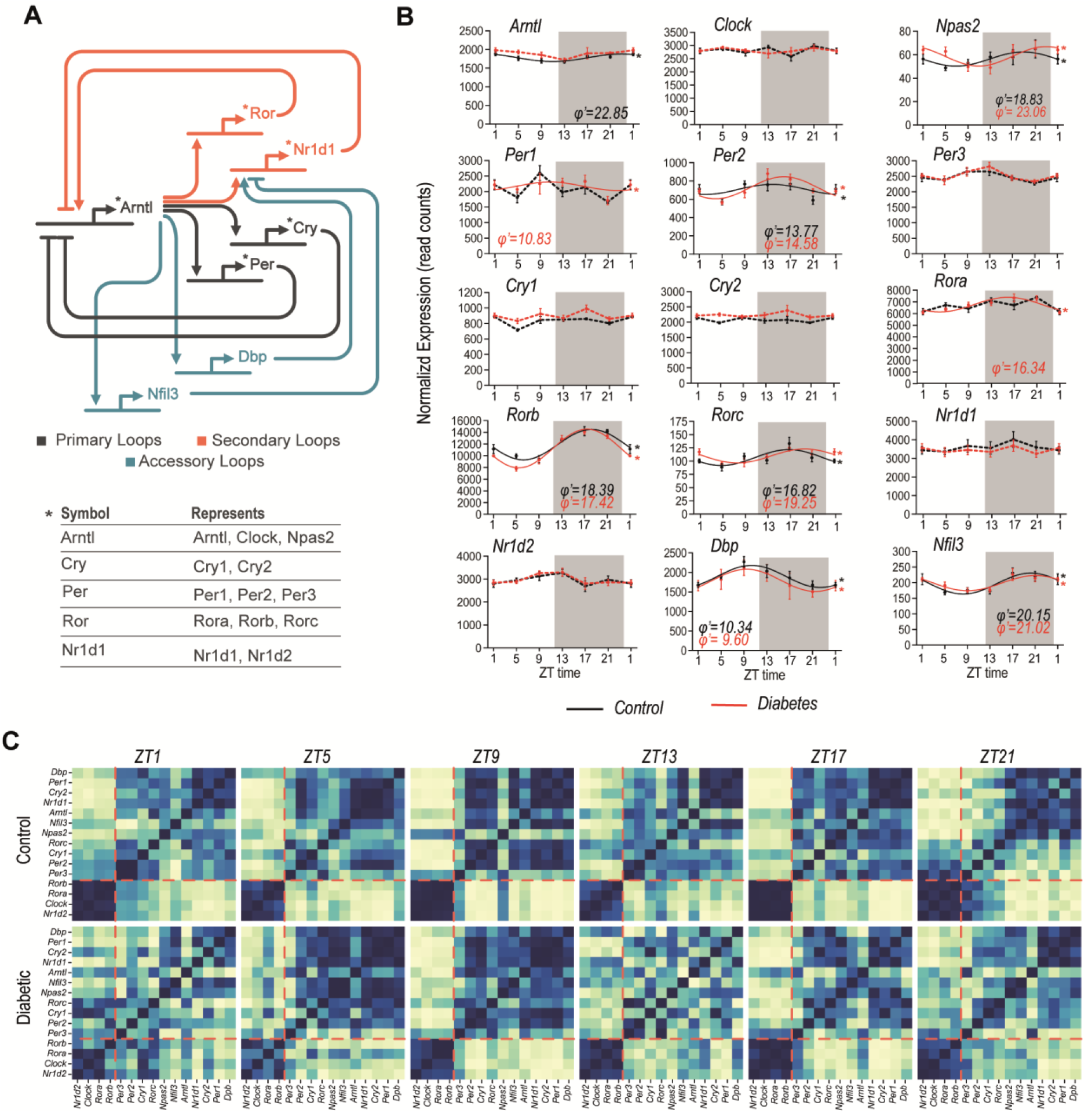
Clock Gene expression in the retina. **(A)** Simplified schematic of the molecular clock, showing some of the main interactions of each gene, including if they inhibit or promote gene expression. For simplicity, genes with similar interactions are represented by a single symbol in the diagram, indicated by the table at the bottom. Primary, secondary, and accessory loops are additionally indicated. **(B)** Expression patterns of clock genes in control (black) and diabetic (red), (mean and standard error for each time-point). Clock genes that have been identified as rhythmic are indicated by an asterisk and solid lines and their respective phase in hours (φ′) beneath. Broken lines indicate no detected rhythmicity. **(C)** Spearman’s rank correlations between each of the 15 clock genes in both control and diabetic groups. Blue indicates strong correlations and yellow/white weakened correlations.

Due to the nature of transcriptional/translational feedback loops, clock gene expression is either directly or indirectly connected and a level of correlation would be expected between genes. For example, when *Arntl* expression is increased the expression of its repressors *Per, Cry, Reverba* should be low. Spearman correlations were calculated between each of the 15 molecular clock genes for each of all the timepoints to reveal if there is disruption in the global expression patterns of all clock genes. While in control retinas, clock gene correlations could be separated into two main groupings, correlations were weakened most prominently at ZT1 and ZT13 in the diabetic retinas (**Fig. 1c**). On closer inspection, correlation disruption can also be identified at other timepoints, as the *Npas2* correlations at ZT9 being strengthened for some genes however weakened for others. Overall, these data indicate a certain level of disruption in the clock machinery in the diabetic retina that might be related to its daily entrainment by the light as these are more evident at dawn and dusk.

### Global daily rhythmic transcriptome in the diabetic retina

Next, we compared the global rhythmic output in control and diabetic mice. 4,700 transcripts in control mice and 5,177 transcripts in diabetic mice were identified as rhythmic using empJTK. When compared to the total number of transcripts identified as expressed (>4 read counts) in the control (19,062) and diabetic retinas (19,122), 25% of expressed transcripts in control and 27% in diabetic retinas displayed rhythmic expression. We then contrasted our own findings to those of previous studies in retinal tissues. Although no available rhythmic RNA-seq datasets exist for the mouse retina, we reanalyzed rhythmic data from microarrays performed in whole-eyes (9) and rhythmic RNA-seq dataset from primate retina (37) using empJTK. However, little overlap existed between the rhythmic transcripts from our study and the available data sets that was not improved with JTK or ASER detection method (data not shown).

In contrast to the subtle effects identified for the clock genes, the level of disruption by diabetes on the global rhythmic transcriptome was significantly greater. Of all rhythmic genes, only 36.7% (2,651) transcripts were shared between control and diabetic retinas, while 28.4% (2,530) gained *de novo* rhythmicity and 35% (2,049) lost diurnal rhythmicity retinas **(Fig. 2A)**. A distinct “gain” or “loss” of rhythmicity can be viewed in the heatmap **(Fig. 2B)**. Pathway enrichment analysis of the genes found rhythmic only in control or only in diabetes or were shared, revealed three major networks being impacted, one related to development and differentiation, another to cell cycle, damage, and metabolism and a third on PI3K-AKT signaling **(Fig. 2B)**. The pathway heatmap **(Fig. 2C)** revealed that most of the pathways were shared, but a lot of them were found to lose rhythmicity in diabetes, such as organ development, PI3-AKT signaling pathway, regulation of cell cycle, response to radiation, cell population proliferation and differentiation, with pathways in cancer and head development being only found to be rhythmic in diabetes **(Fig. 2D)**. E2f1, Sox10 and SP1 were found to likely regulate the expression of genes found only in diabetes **(Fig 2E)**.

**Figure 2.**
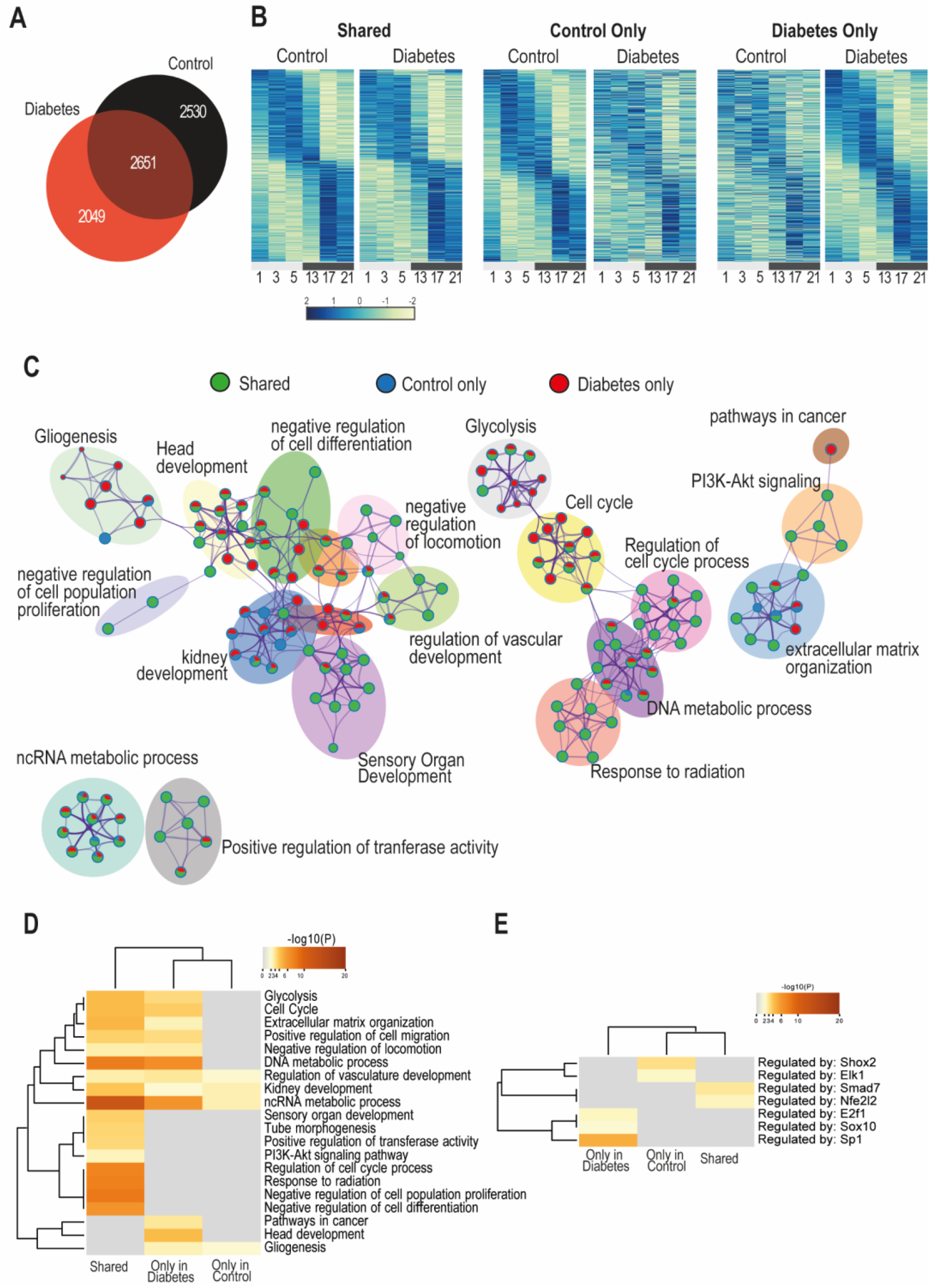
Disorganization of rhythmic transcripts in diabetes. **(A)** Venn diagram showing the overlap between transcripts identified as rhythmic in the control and diabetic retinas. **(B)** Heatmaps indicate loss, gain or retention of rhythmicity in control and diabetic retinas. Transcripts expression were min/max normalized and sorted by harmonic regression predicted acrophase. Each row represents a gene identified as cycling in each control or diabetic or both. Each column represents a single time-point, where biological replicates expression was averaged. Blue to yellow hue represents gene expression from max to min. **(C)** Metascape network pathway analysis of transcripts that were identified as rhythmic. Nodes are coded with colors depending on if they represent rhythmic genes found only in control (blue), only in diabetic (red), or shared in both control and diabetic retinas (green). **(D)** Gene set enrichment analysis revealing the pathways identified to be rhythmic only in diabetes, only in control or shared. **(E)** Predicted regulators for the rhythmic genes in each category.

Plotting the time of the acrophases, determined by harmonic regression, revealed two clear peaks in transcription in both control and diabetic retinas, one during the day and one during the night, separated by an approximately 12-hour difference (**Fig. 3A-B**). In control retinas, most transcripts peaked between ZT6 and ZT8, and ZT18 and ZT20 (**Fig 3A**). However, in the diabetic retinas although the 12-hour expression axis was retained the transcripts were slightly phase advanced, between ZT5 and ZT7 and ZT17 and ZT19 (**Fig 3B**). In addition, the two peaks in the diabetic group are more widely distributed, with neither as prominent as those in control retinas. These patterns were true for both coding and non-coding transcripts.

**Figure 3.**
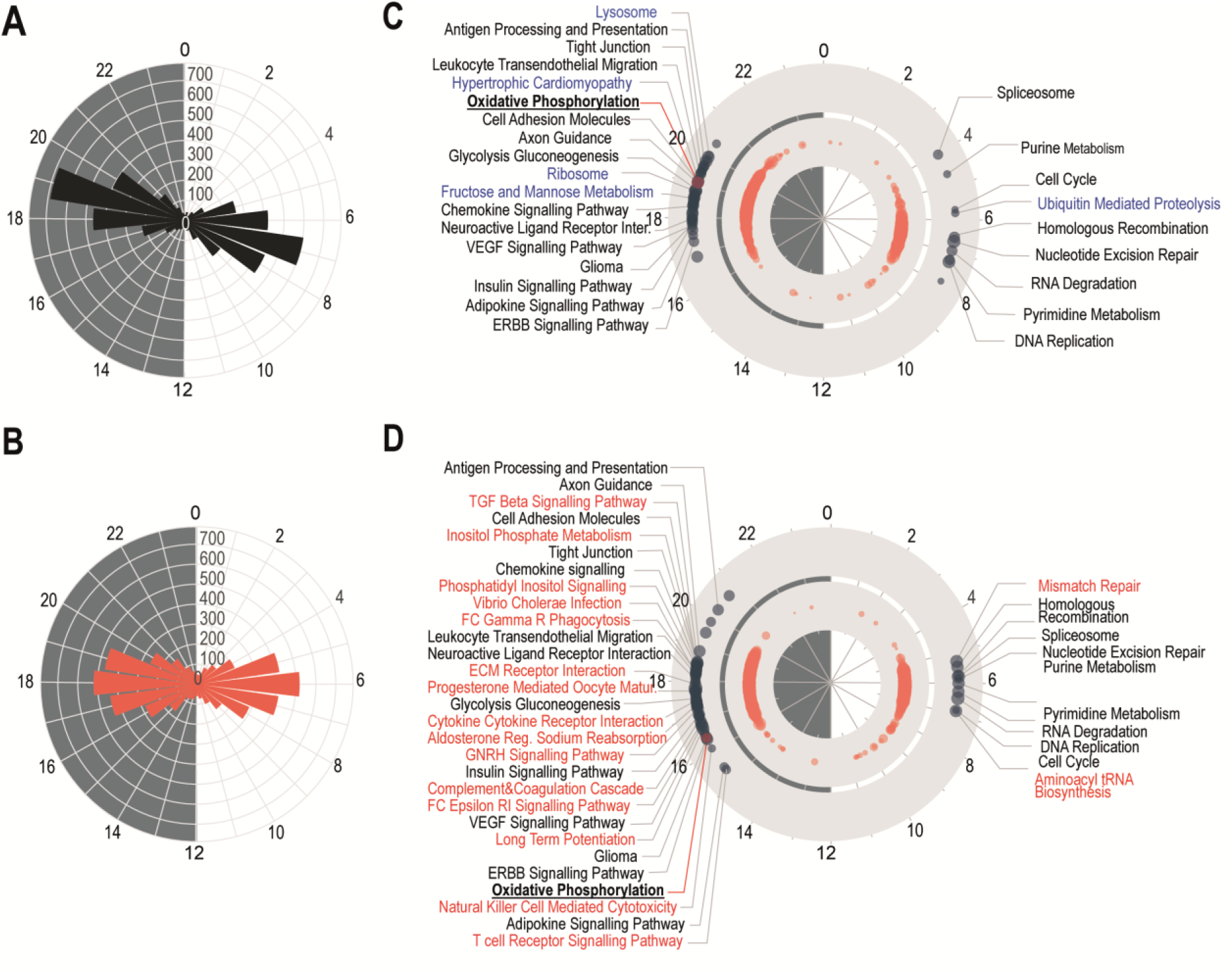
Phase differences in rhythmic transcripts. **(A-B)** Radial plots of the distribution of the peak phase of rhythmic transcripts in control **(A)**, and in diabetic **(B)** retinas. **(C-D)** Phase distribution over the 24h cycle of representative KEGG pathways in control **(C)** and diabetic **(D)** retinas. Phases were calculated using Set Enrichment Analysis (PSEA) for rhythmic transcripts. KEGG pathways which were uniquely identified as rhythmic in control are highlighted in blue and in diabetes in red.

We then conducted a functional analysis whilst considering the acrophases of rhythmic transcripts (34). This identified that the peak of expression during the day and during the night have separate roles in the retina **(Fig3 C-D)**. Genes peaking at the mid-day were enriched in pathways related to cell cycle, DNA replication and RNA degradation, while those that peaking at mid-night were enriched for metabolism, inflammation and growth factor signaling pathways, indicating a clear functional compartmentalization of the retinal transcriptome between day and night. Interestingly, there were fewer enriched KEGG pathways during the day when compared to the night. Most importantly, we identified unique KEGG pathways which were enriched in either the control or the diabetic retina. During the day, control retinas (**Fig 3C**) showed enrichment of the ubiquitin-mediated proteolysis pathway genes which was lost in diabetes. In contrast, diabetic retinas have enriched pathways related to mismatched repair and tRNA biosynthesis pathways not found to be rhythmic in control retinas (**Fig 3D**). During the night, more pathways were altered by diabetes, especially those related to inflammation, hormonal and cytokine signaling, while a few were absent, such as ribosome, fructose and mannose metabolism, lysosome and pathways related to cardiomyopathy (**Fig 3D**). The genes in some pathways exhibited an even earlier peak phase in diabetes than the total cycling transcripts. Like the earlier peak phase of the total cycling transcripts, KEGG pathways also exhibited an earlier peak in diabetic retina compared to control, but some more than others. Oxidative phosphorylation, for example, was advanced by almost 4 hours in diabetes compared to control.

### Amplitude and phase changes in the rhythmic transcriptome in the diabetic retina

We first aimed to identify whether diabetes resulted in differential expression at individual timepoints (ZT1, -5, -9, -13, -17, -21) or among all samples in control and diabetic groups independent of the time, but we only identified few genes been significant different with these approaches (**Sup. Table 5**). The very low numbers of differentially expressed transcripts perhaps indicates there were no differences in the relative expression of each transcript, although they may display changes in rhythmicity patterns.

To further examine the differential rhythmic expression of genes between control and diabetic retinas, we used the differential rhythmicity analysis, DODR (36), and identified a total of 478 genes as differentially rhythmic. We split these into 2 main groups: transcripts that displayed an altered amplitude (**Fig. 4A**), and transcripts that displayed a phase shift >1-hour (**Fig. 4B**). A higher number of genes identified as phase-shifted (405 transcripts) compared to genes with altered amplitude (154 transcripts). This was also confirmed using cosinor analysis where indeed most of the genes showed a phase-shift in their expression rather than an amplitude change (data not shown). Most shifted genes were phase-advanced in the diabetic retinas.

**Figure 4.**
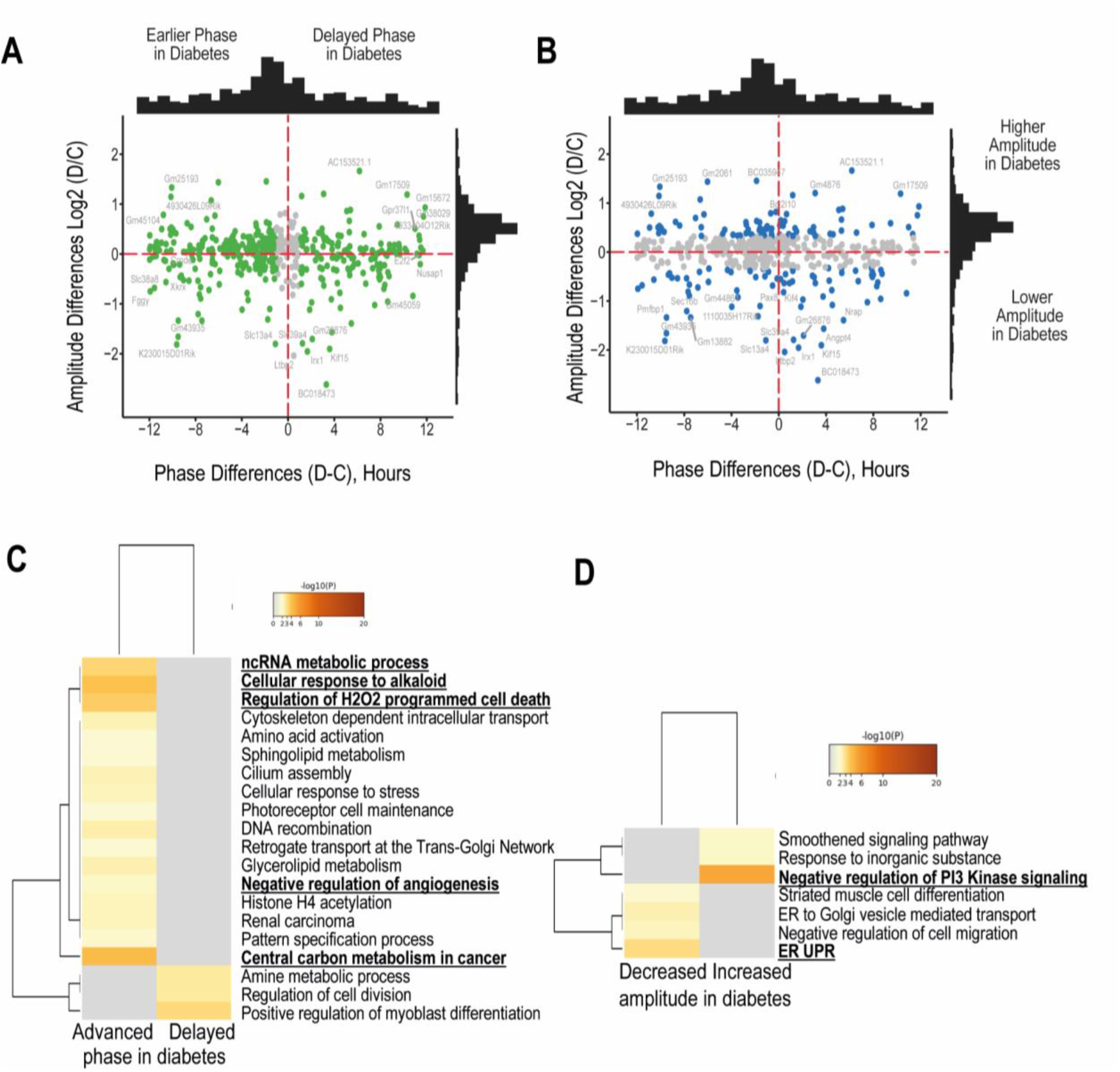
Phase and amplitude changes of differentially rhythmic transcripts between controls and diabetic retinas. DORD analysis was performed to identify differential rhythmic transcripts based on amplitude and acrophase. **(A)** All transcripts that showed a statistically significant difference in phase are highlighted in green with the distribution of the number shown on the top. **(B)** All transcripts with differences in amplitude are shown in blue. A distribution of the number of genes shown on the right. **(C)** Differentially rhythmic genes were further subdivided into those that displayed advanced or delayed phase and Metascape analysis was performed for pathway and enrichment analysis; **(D)** Metascape analysis for genes that had increased or decreased amplitude in diabetes.

Enrichment analysis of the differentially rhythmic genes (**Fig. 4C**) revealed that central carbon metabolism, ncRNA-regulated metabolic processes, cellular response to alkaloids and oxidative induced cell death were the most enriched pathways, while circadian control was not identified to be differentially rhythmic in the analyzed retinas. Sphingolipid metabolism and histone H4 acetylation were also among the pathways that were enriched in genes with advanced phase in diabetes. Genes that showed delayed phase were enriched in cell division and differentiation pathways. In contrast, pathways that exhibited increased amplitude with diabetes (**Fig 4D**) were enriched in the regulation of PI3K signaling, and those that had reduced amplitude, in endoplasmic reticulum stress and the unfolded protein response (UPR).

Canonical pathway analysis of all the genes that were identified to be differentially rhythmic in control and diabetes included phototransduction, glucose metabolism, HIF1a signaling, ER stress, DNA damage and cell cycle checkpoints to be affected (**Fig 5A**). Upstream regulator analysis suggested that COMMD1, HIF1A, EGLN, PIK3R1 and MTORC1 as major upstream regulators for the phase shifts, while HSP90B1, YWHAZ, CNGA3, COL2A1 and TFRC (**Fig 5B**) as the major upstream regulators for the amplitude changes observed in the diabetic retina.

**Figure 5.**
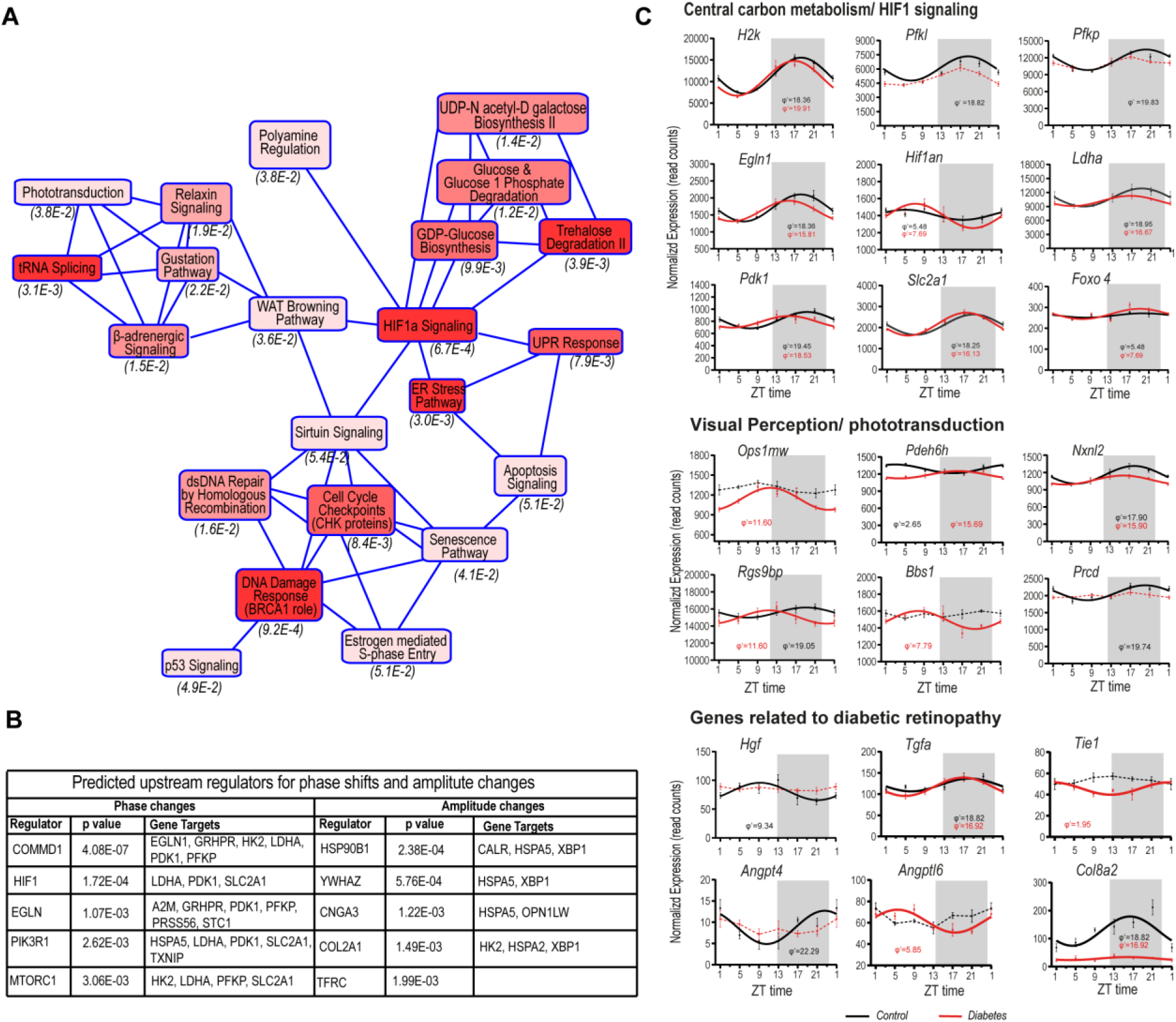
Pathway analysis of differential rhythmic genes. **(A)** Graphic summary of canonical pathway analysis with IPA Core Analysis illustrating how enriched canonical pathways are related to one another. **(B)** Upstream Regulator Analysis indicating transcriptional regulators likely to be related to phase and amplitude changes. **(C)** Expression patterns of genes that showed different rhythmicity in diabetic retinas compared to control, involved in central carbon metabolism and HIF1 signaling, visual perception/phototransduction and pathogenesis of DR.

We then explored some of the pathways that were identified to be driving the phase shifts. Among the genes that were differentially regulated in the central carbon metabolism and Hif1a signaling, *Hk2, Pfkl, Pfkp, Egln1, Ldha, Pdk1, Glut1* were advanced by few hours as shown in **Fig 5C**. However, genes related to photoreceptors were significantly phase shifted with *Pdhe6* and *Rgs9bp* exhibiting around 10-hour phase-shifts. This altered rhythmicity involves some genes associated with DR, such as *Tgfa, angpt4, angptl6, hgf* and *Col8a2*. Overall, these data indicate that a possible metabolic dysfunction involving metabolism and visual pathways around the daily cycle drives the phase shifts observed in the diabetic retina.

## Discussion

Emerging evidence for circadian disruption in the diabetic retina (14, 15, 38, 39) prompted us to a detailed transcriptomics analysis in the retina of T1D mice with focus on real-life conditions (light/dark) rhythmicity. We used deep mRNA sequencing to examine the effects of established type 1 diabetes in the daily rhythmic output of the retinal transcriptome and showed a significant misalignment of the rhythms of retinal gene expression that takes place early after the onset of diabetes and before the appearance of DR pathology. We confirmed that a 12-hour axis in transcriptional compartmentalization between day and night was also present in the retina, as reported for other organs (37) and it remained in diabetes but is phase-shifted by 1-3 hrs, or up to 10 hrs or more for some individual genes. An advancement of phase rather than delay or changes in amplitude were mostly observed in the differentially rhythmic genes. Interestingly, these phase shifts seem to be mainly driven by the phase advance observed in pathways related to metabolic dysfunction and visual pathways involving HIF1a and mTOR1 signaling pathways, rather than alterations in the phase of the circadian retinal clock, suggesting that in early diabetes a metabolic jet lag occurs in the retina.

While it is established that diabetes distorts clock gene expression in the retina (12, 15, 38), the current study detected only subtle changes in the amplitude and the phase of expression of clock genes. Clock genes remained in phase, except for Npas2, but some lost their daily rhythmicity (*Rora, Rorc* and *Nr1d1*) while others gained rhythmicity (*Per2*). While this is unexpected under light/dark conditions, a degree of discoordination of clock gene relationships were observed especially at dawn and dusk, indicating subtle effects of diabetes in the synchronization of the clock machinery to light. While our data suggest that clock gene expression remained in phase, we do not exclude that specific cell clocks are influenced by diabetes, nor that the subtle changes are due to loss of specific cell populations within the diabetic retina or alterations of clock regulation within the retina. We used bulk mRNA sequencing of the whole retina, which reflects the average gene expression signature of the diverse cell populations within the retina. This approach can mask changes in clock gene expression in specific cell-types that have proportionately low representation within the retina, such as endothelial cells. Moreover, it is well established that the retina, a highly heterogeneous and laminated tissue, harbors several clocks distributed throughout the various cell types and between retinal layers, each with different phases and period lengths (40, 41). This may limit the utility of bulk mRNA sequencing techniques to dissect how a disease influences clock regulation in rare populations. Nevertheless, our study showed that, in this early time point and under entrainment conditions, clock gene expression was found in phase, even if they exhibited subtle changes in amplitude which could be driven by cell death, or impact on the clock regulation itself.

Very few studies have analyzed the diurnal or circadian transcriptome in the murine retina (9), primates (37), drosophila (42) and zebrafish(43) none of which were done in the context of diabetes. To our knowledge, the current study is the first that maps the effects of diabetes on the retina’s rhythmic transcriptome around the daily cycle. A previous study examined the effects of diabetes on extraorbital lacrimal glands using the same approaches (44) and identified a phase advance due to diabetes in the diurnal transcriptome, in agreement with our finding in the retina. In addition, they identified rhythmic genes with only little overlap between controls and diabetes, indicating a major reorganization of the daily rhythmic transcriptome by diabetes. Similarly, Vancura et al. (45) performed a microarray in the retina of T2D mice (db/db) comparing two time points (one at day and one at night). The authors identified 31 differentially expressed genes between day and night in diabetes and 278 in control, suggesting a disruption of the rhythmic output in the diabetic retina with a small overlap of the common genes that change from day to night. We identified that there were more genes gaining rhythmicity in diabetes compared to control, similarly with Jiao et al. data (44). Genes that “gained de novo” rhythmicity due to diabetes belonged to pathways of responses to growth factors, transmembrane receptor protein kinase signaling, negative regulation of cell proliferation and cellular component organization, positive regulation of cell death and head development. Among those genes, growth factors such as *Vegf, Tgf-β1, Pdgf, Hgf* and *Igf-1* have been associated with DR and are actively investigated for therapeutic targets for treating aberrant angiogenesis. The gaining of rhythmicity may not be driven by the circadian clock itself, but it does reveal a topological preference in the temporal timeline of a day, bringing into focus the application of chronotherapies for the management of DR as most of the genes peaking during the night and therefore gaining rhythmicity are active targets for the management of DR.

Our analysis revealed that the retina exhibits a 12 hr transcriptional compartmentalization, which is induced by light and fine-tuned by the circadian clock. This transcriptional axis has been previously observed in other organs in primates (37) but not the retina. We report here that day peaking genes favored DNA repair, RNA splicing and ribosomal protein synthesis, whereas night peaking genes are related to metabolic processes and growth factor signaling and the major therapeutic targets for DR. Although the 12 hr transcriptional axis is retained in the diabetic retina, it was phase advanced by approximately 1-3 hour with a wider distribution. Interestingly, we found more alterations in the phase than in the amplitude of the rhythms, shown in other ocular tissues as well (44). This can render the retina more susceptible to certain challenges, such as its response to light damage. Moreover, even when small phase misalignments with the daily cycle are imposed, the damage to the tissue may accumulate when these conditions become chronic. For example, a chronic small phase misalignment of the internal clock with the external light/dark cycle can have severe impact on the metabolic and cardiovascular health of mice (46). Although we suggest that this may not be driven by the circadian clock under light/dark conditions, a study in a type 2 diabetic Per2::luciferase mouse showed that the period of the circadian peripheral metabolic clocks such as the liver and white adipose tissue is significantly reduced (and therefore phase is advanced) (47), so additional studies need to be conducted to understand what could drive these phase shifts.

Our findings led us to propose a possible mechanism that could facilitate the damage occurring in the diabetic retina. The hypothesis is described in **Figure 6** and postulates that diabetes creates a metabolic jet lag in the retina driven by a phase advancement of pathways related to hypoxia and metabolism, while the circadian clock rhythms remain entrained to the light/dark cycle, driving clock-controlled responses in phase with the environment. The implications of these phase shifts are not yet known, but we hypothesize that the chronic internal misalignment, observed in the early stages of diabetes, contributes to the pathology of DR by increasing oxidative stress and neuronal damage early on. This hypothesis postulates that having a functional entrainable retinal clock in diabetes could have a pathogenic role for the progression of DR and predicts that when this internal jet lag is corrected by, for example, deletion of the retina circadian clock with retina specific knockout of Bmal-1 or with correction of the metabolic phase shifts, then the progression of DR could be slowed. Future studies will be required to confirm this.

**Figure 6.**
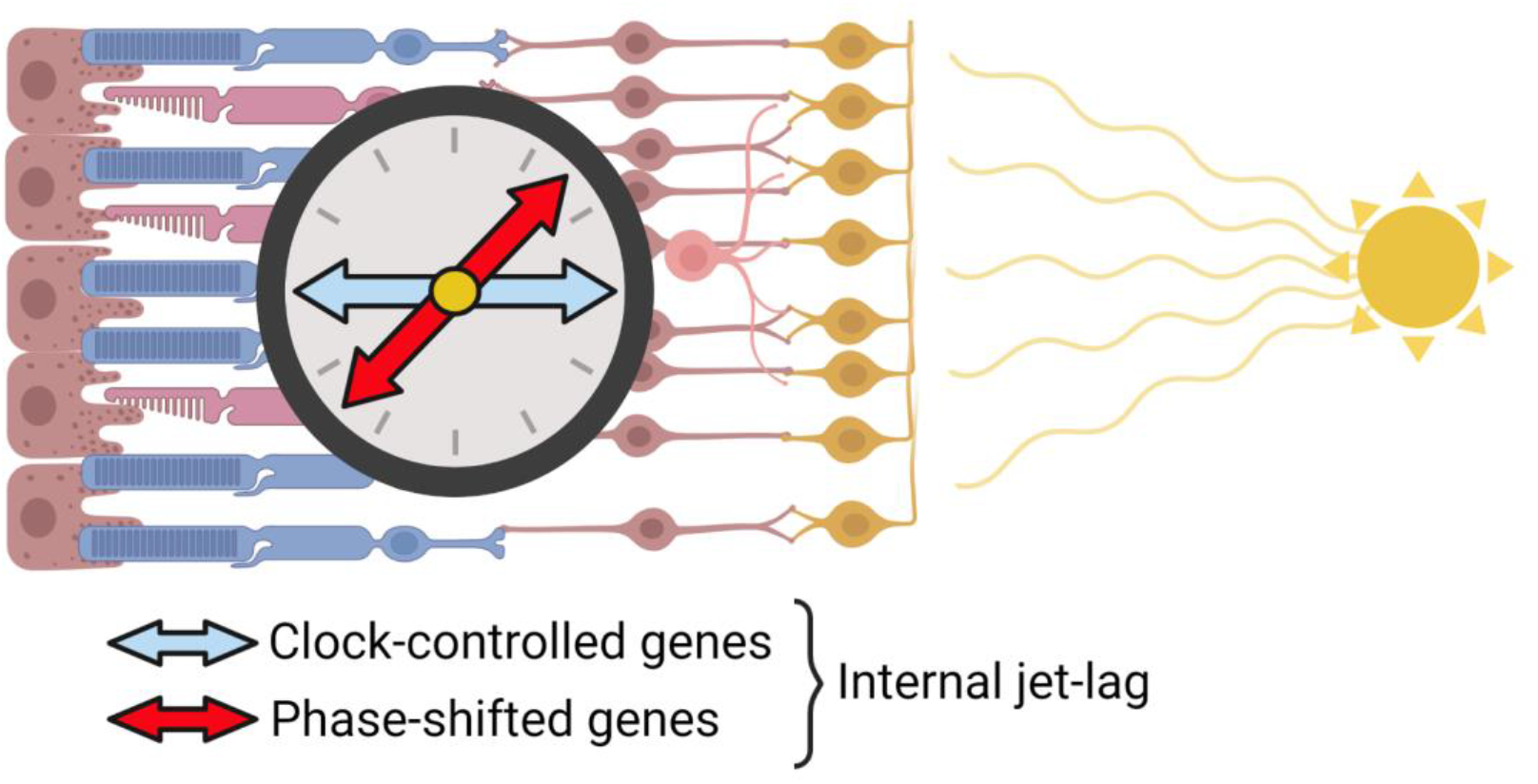
Proposed model for metabolic jet-lag in the diabetic retina. In early diabetes, the retinal clock is entrained to the light/dark cycle and drives clock-controlled output at certain times of the day. However, some metabolic and genes related to visual perception exhibit a phase advance characterized from 2-10 hours which may create an internal jet lag and breakdown of the coordination between metabolic and circadian rhythms within the retina. This phase advance is likely driven by major transcription factors regulating the hypoxic and metabolic response. Created with BioRender.com.

Indeed, a pathological role of Bmal1 in the progression of DR was recently proposed by Vancura et al. (45), who showed that Bmal1 deletion spared the retina from the effects of diabetes. Moreover, recent work from Jidigan et al. (48) showed that the neuronal clock in the context of pathological neovascularization is detrimental and its deletion led to protection from aberrant angiogenesis. Disease-mediated disruption of circadian rhythms therefore may emerge as an important factor in the disease progression. This study suggests that hypoxia occurring within the diabetic retina could drive a circadian dysfunction promoting the development of DR.

## Acknowledgments

We would like to thank the Genomics CTU at QUB and colleagues that provided insightful conversations, Thomas Friedel for his technical support. Finally, we would like to thank the funding sources: JDRF grant 1-FAC-2019-878-A-N (to EB) and NIH grants R01 DK093954 and DK127308, U01DK127786, UC4 DK 104166, 2-SRA-2019-834-S-B, JDRF 2-SRA-2018-493-A-B (to CEM).

## Author Contributions

EB designed experiments; RPS, VKT, DAS and EB designed, performed and discussed bioinformatics analysis; RPS wrote the manuscript; RPS, EB, HRW, ODB, AWS discussed and interpreted the results; RPS, EB, HRW ODB, DAS reviewed and revised; CEM, DAS, AWS provided edits to the manuscript.

## Notes

### Competing Interest Statement

The authors have declared no competing interest.

