## Supplementary material for "Mapping the daily rhythmic transcriptome in the diabetic retina": Supl. Figures

**Supplemental Material:**

**Supplemental Figure 1| Experimental design for sample collection.**

Ins2 Akita mice and their littermate controls were bred and kept until Mice were kept in 12hr:12hr Light:Dark (LD) conditions throughout. Samples were collected at 4-hour intervals throughout a 24-hour period starting at ZT1 and ending at ZT21. 4-5 biological replicates were taken for each experimental group at each timepoint.

**Supplemental Figure 2 | Patterns of rhythmicity in identified known clock-controlled genes**

Transcripts per million in known clock-controlled genes in the retina of control and diabetic mice. Expression patterns of clock genes in control (black) and diabetic (red), also showing mean and standard error for each time-point. Clock genes that have been identified as cycling are indicated by an asterisk and solid lines and display the phase in hours (φ′) beneath. Broken lines indicate no detected rhythmicity. n= (4-5)/ time point.

**Supplemental Figure 1**

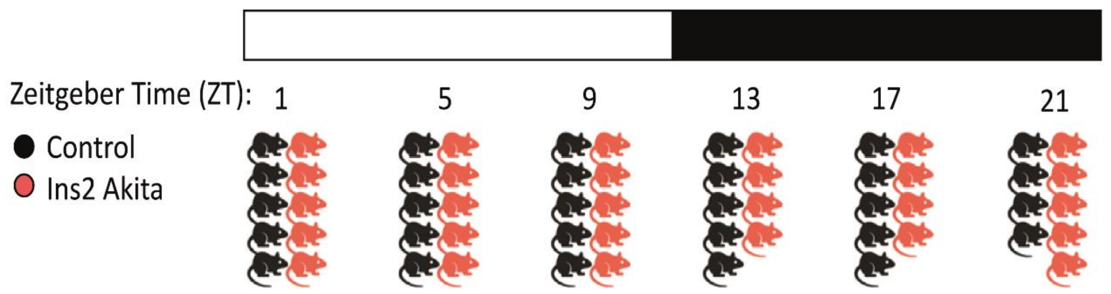


**Supplemental Figure 2**

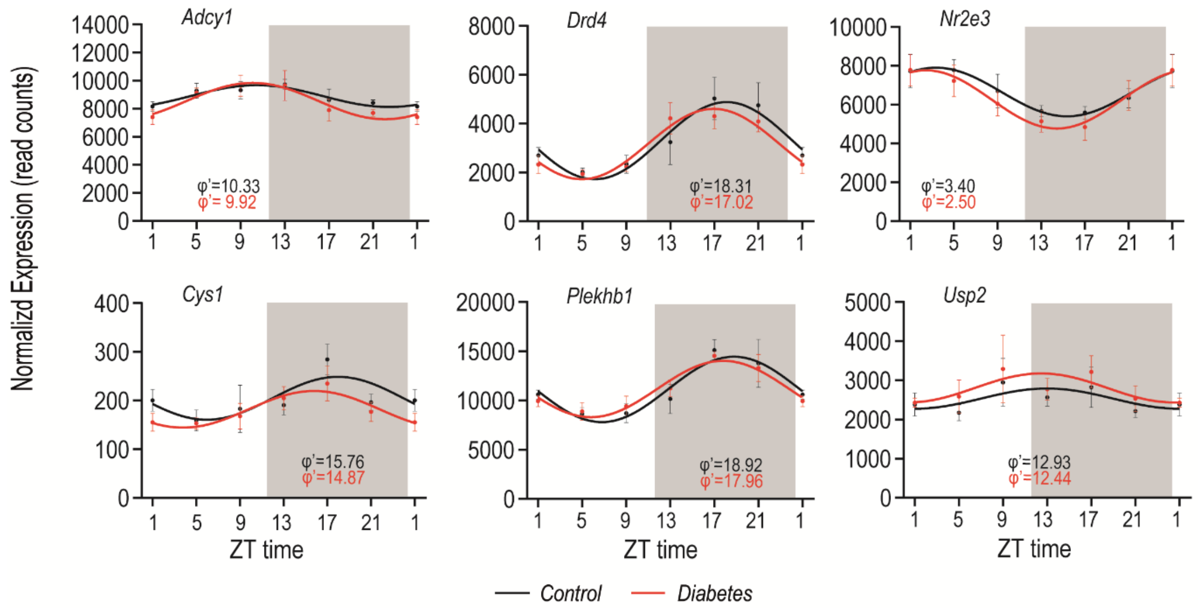
